# Broadband Dynamics Rather than Frequency-Specific Rhythms Underlie Prediction Error in the Primate Auditory Cortex

**DOI:** 10.1101/821942

**Authors:** Andrés Canales-Johnson, Ana Filipa Teixeira Borges, Misako Komatsu, Naotaka Fujii, Johannes J. Fahrenfort, Kai J. Miller, Valdas Noreika

**Author notes:** Correspondence (A.C.-J.), (V.N.). These authors contributed equally to this work (Andrés Canales-Johnson, Ana Filipa Teixeira Borges).

## Abstract

Detection of statistical irregularities, measured as a prediction error response, is fundamental to the perceptual monitoring of the environment. We studied whether prediction error response is associated with neural oscillations or asynchronous broadband activity. Electrocorticography (ECoG) was carried out in three male monkeys, who passively listened to the auditory roving oddball stimuli. Local field potentials (LFP) recorded over the auditory cortex underwent spectral principal component analysis, which decoupled broadband and rhythmic components of the LFP signal. We found that the broadband component captured the prediction error response, whereas none of the rhythmic components were associated with statistical irregularities of sounds. The broadband component displayed more stochastic, asymmetrical multifractal properties than the rhythmic components, which revealed more self-similar dynamics. We thus conclude that the prediction error response is captured by neuronal populations generating asynchronous broadband activity, defined by irregular dynamical states which, unlike oscillatory rhythms, appear to enable the neural representation of auditory prediction error response.

**Significance Statement:** This study aimed to examine the contribution of oscillatory and asynchronous components of auditory local field potentials in the generation of prediction error responses to sensory irregularities, as this has not been directly addressed in the previous studies. Here, we show that mismatch negativity – an auditory prediction error response – is driven by the asynchronous broadband component of potentials recorded in the auditory cortex. This finding highlights the importance of non-oscillatory neural processes in the predictive monitoring of the environment. At a more general level, the study demonstrates that stochastic neural processes, which are often disregarded as neural noise, do have a functional role in the processing of sensory information.

## Introduction

Detection of novel sensory information enables adaptive interaction with the surrounding environment (Clark, 2013; Whitmire and Stanley, 2016). In the predictive coding framework of brain functioning, this interaction is characterized by a reciprocal loop between sensory predictions and prediction error signals (Bastos et al., 2012; Friston and Kiebel, 2009). Neural mechanisms of prediction error are typically studied by presenting a series of “standard” stimuli with intermittently occurring deviant stimuli, also called “oddballs”, and by contrasting brain responses between these stimulus categories (Chennu et al., 2013; Lumaca et al., 2019; Parras et al., 2017). This way, event-related potentials (ERP) and a range of neural oscillations have been identified as neural markers of prediction error. The most widely studied deviance ERP is the auditory mismatch negativity (MMN) – a negative deflection of electrical event-related potential recorded on the scalp or using intracranial electrodes (Halgren et al., 1995; Näätänen et al., 1978; 2007; Rosburg et al., 2005). MMN originates from the primary auditory cortex (Alain et al., 1998; Alho, 1995; Edwards et al., 2005), and it peaks around 150-200 ms in humans, whilst the peak latencies below 100 ms are typically reported in monkeys (Camalier et al., 2019; Javitt et al., 1992; Komatsu et al., 2015). In addition to MMN, prediction error responses are observed in a variety of frequency ranges including theta (3-8 Hz) (Choi et al., 2013; Fuentemilla et al., 2008; Hsiao et al., 2009; Ko et al., 2012; MacLean et al., 2014), alpha (8-12 Hz) (Ko et al., 2012; MacLean et al., 2014), beta (14-30 Hz) (Haenschel et al., 2000; MacLean et al., 2014) and gamma (>30 Hz) (Dürschmid et al., 2016; Edwards et al., 2005; Eliades et al., 2014; Haenschel et al., 2000; MacLean et al., 2014; Marshall et al., 1996) ranges.

Several interpretations could be formulated aiming to explain the abundance of prediction error responses in the frequency dimension. First of all, there could be multiple independent neural mechanisms sensitive to stimulus deviance. This suggestion, however, does not explain why there would be so many distinct mechanisms with an identical functional role. Alternatively, frequency-specific detectors of prediction error might be only partially independent, forming hierarchical cross-frequency interactions. For instance, rhythms of different frequency bands could drive each other, e.g. delta phase could modulate theta amplitude and theta phase could modulate gamma amplitude in the auditory cortex (Lakatos et al., 2005). Yet another possibility – which we pursue in the present study – is that a broad frequency range of deviance responses, including theta, alpha, beta and gamma bands, points to a *broadband* prediction error response, which is not restricted to a particular frequency band, but instead is driven by an arrhythmic or asynchronous neural signal across a wide frequency range. In fact, a large number of studies reported deviance effects to run across several frequency bands (Chao et al., 2018; Haenschel et al., 2000; Hsiao et al., 2009; Ko et al., 2012; MacLean et al., 2014), arguably alluding to arrhythmic processing of unexpected stimuli.

The electrophysiological signal recorded by scalp EEG or local field potentials (LFP) is a summed activity of both postsynaptic and action potentials. Post-synaptic potentials contribute to the rhythmic oscillations of different frequency bands (Buzsaki et al., 2012), reflecting neural synchrony at specific timescales. Contrary to this, empirical data analysis and modeling suggest that the average neuronal firing rate produces asynchronous, broadband changes across a wide frequency range (Buzsáki et al., 2012; Guyon et al., 2021; Henrie and Shapley, 2005; Hwang and Andersen, 2011; Li et al., 2019; Medvedev and Kanwal, 2004; Mukamel et al., 2005). Such rhythmic and broadband components of LFP signal can be decomposed using spectral principal component analysis (spectral PCA) (Miller et al., 2009a, 2009b, 2017; Miller, 2019), this way separating synchronous and asynchronous neural activity. The broadband component of the LFP power spectrum is commonly characterized by a power-law function (Freeman and Zhai, 2009; He, 2014; Hermes et al., 2019), which reflects the lack of any specific temporal beat (e.g. 10 Hz) in the signal. Contrary to this, rhythmic components produce frequency-specific spectral peaks that deviate from the power law. In fact, the electrocorticography power is characterized by at least three different power-law regions of which the transitions vary across individuals and recordings in humans (Chaudhuri et al., 2017; He et al., 2010) and non-human primates. The functional relevance of this heterogeneous scaling is discernible as, for instance, levels of arousal across a gradual progression from awake to anesthesia (Gifani et al., 2007) or deep sleep (Ma et al., 2006; Weiss et al., 2009) can manifest selectively within power-law changes at different timescales. Such complex dynamics across different LFP timescales can be characterized by multiscale multifractal analysis (MMA; Gierałtowski et al., 2012), developed to analyze signal fluctuations on a wide range of timescales like those observed in LFP signals.

In the present study, we aimed to assess whether such broadband neural dynamics rather than frequency-specific rhythms underlie prediction error in the auditory cortex in the primate brain. Furthermore, we sought to contrast the *multiscale dynamics* of the broadband LFP component to that of the rhythmic components. Importantly, while several previous LFP and EEG studies have linked MMN to gamma range activity (Dürschmid et al., 2016; Edwards et al., 2005; Eliades et al., 2014; Haenschel et al., 2000; MacLean et al., 2014; Marshall et al., 1996), and often referred to it as a “broadband” neural signal, spectral power of raw signal conflates genuinely asynchronous broadband activity with gamma oscillatory activity (for a difference in visual processing, see Hermes et al., 2019). Thus, it remains uncertain whether the reported gamma correlates of MMN are driven by gamma oscillations or non-oscillatory broadband activity. Addressing this issue, we used spectral PCA analysis which decoupled broadband activity from the oscillatory components of the neural signal and enabled us to assess their distinct functional associations with MMN.

Our key hypotheses were based on the predictive coding explanation for the mismatch negativity – or indeed any violation or omission response. In brief, in predictive coding formulations there are two neurocomputational mechanisms in play. First, mismatches between ascending sensory afferents and descending predictions are registered by prediction error units (usually considered to be superficial pyramidal cells) (Bastos et al., 2012; Shipp, 2016). These prediction error responses are then passed to higher hierarchical levels of processing to drive Bayesian belief updating, optimize descending predictions, and resolve prediction errors – in a continuous process of recurrent hierarchical message passing. According to this framework, a change in the predicted frequency would be reported by prediction error units tuned to both the new and old (i.e., oddball and standard) frequencies. Physiologically, these responses are mediated by changes in neuronal firing rates; either over broad frequency bands (e.g., as measured by multiunit activity) or, on some accounts, high gamma frequencies (Bastos et al., 2015a; Bastos et al., 2015b).

The second neurocomputational mechanism of predictive coding corresponds to an optimization of the gain or excitability of prediction error populations, immediately following an unpredicted sensory input. Computationally, this corresponds to increasing the precision of prediction errors – so that they have a greater influence on belief updating (Clark, 2013; Feldman and Friston, 2010). In engineering, this corresponds to a change in the Kalman gain (Rao and Ballard, 1999). Physiologically, this is manifest in terms of an increased postsynaptic gain or loss of self-inhibition. Electrophysiologically, this change in inhibition-excitation balance would be expressed in terms of a (deviant induced) dynamical instability, of the sort characterized by scale-invariance. Please see Friston et al. (2012) for a mathematical analysis in terms of Lyapunov and scaling exponents (including Fig. 7 for a detailed simulation of induced auditory responses).

On the predictive coding account, the first mechanism rests upon top-down predictions based upon a generative model of the auditory stream. This provides a formal description of the model adjustment hypothesis (Garrido et al., 2009). The second mechanism depends upon changing postsynaptic sensitivity, which offers a formal explanation for sensory-specific adaptation and related phenomena (e.g., spike-rate adaptation). From our perspective, both mechanisms make a clear prediction: responses to deviant stimuli will be expressed in broadband (non-oscillatory) response – reflecting the activity of prediction error populations. Furthermore, such broadband signals should show evidence of increasing dynamical instability compared to the oscillatory responses as measured in terms of scaling exponents. Based on the current knowledge reviewed above about the two mechanisms of predictive coding, we make the following predictions: 1) auditory oddball responses to deviant stimuli will be expressed in broadband (non-oscillatory) component of electrophysiological/ECoG signal – reflecting the activity of prediction error neural populations; 2) these broadband responses will show increased dynamical instability relative to that of frequency-specific rhythms, as measured in terms of scaling exponents at several timescales. We tested these two hypotheses using PCA-based spectral decoupling PCA and multiscale multifractal analysis. Regarding broadband MMN response, while predictive processing entails both *local* predictions across adjacent stimuli as well as *global* predictions across different sequences of stimuli (Chennu et al., 2013, 2016), we focused on the relatively fast local prediction errors in the present study.

## Methods

### Subjects

We tested three adult male common marmosets (*Callithrix jacchus*) that weighed 320–380 g. Monkeys were implanted with an ECoG electrode array under general anesthesia, and all efforts were made to minimize suffering. All surgical and experimental procedures were performed under the National Institutes of Health Guidelines for the Care and Use of Laboratory Animals and approved by the RIKEN Ethical Committee (No. H26-2-202). ERP data of one monkey (Fr) was reported previously (Komatsu et al., 2015), whereas datasets of monkeys Go and Kr are new.

### Implantation of ECoG arrays

Chronically implanted, customized multichannel ECoG electrode arrays (Fig. 1B–E) (Cir-Tech Inc., Japan) were used for neural recordings (Komatsu et al., 2015; 2017). We implanted 32 (the left hemisphere of monkey Fr), 64 (the right hemisphere of monkey Go), and 64 (the right hemisphere of monkey Kr) electrodes in the epidural space. For the 32 electrode array, each electrode contact was 1 mm in diameter and had an inter-electrode distance of 2.5–5.0 mm (Komatsu et al., 2015). For the 64 electrode array, each electrode contact was 0.6 mm in diameter and had an inter-electrode distance of 1.4 mm in a bipolar pair (Komatsu et al., 2017). The electrode-array covered the frontal, parietal, temporal, and occipital lobes. The additional 4 electrodes of monkey Fr covered part of the right frontal lobe. The animals were initially sedated with butorphanol (0.2 mg/kg i.m.), and surgical anesthesia was achieved with ketamine (30 mg/kg i.m.) and medetomidine (350 μg/kg i.m.). The animals were then positioned in a stereotaxic frame (Narishige, Japan) and placed on a heating pad during surgery. Vital signs were monitored throughout surgery. Implantation of the electrode-arrays involved the removal of a bone flap (~2 cm along the anterior-posterior axis and ~1 cm along the mediolateral axis) over the parietal cortex. The array was advanced into the epidural space. After positioning the electrode-array, connectors were attached to the bone using dental acrylic and titanium (size 1.0 × 0.1mm) or PEEK (size 1.4 × 2.5 mm) screws. The reference electrodes were placed in the epidural space and the ground electrodes in the episkull space. The anti-inflammatory corticosteroid dexamethasone (1.25mg/kg, i.m.) was administered after surgery to prevent brain swelling. The animals were given antibiotics and analgesics daily for 5 days after surgery. Following the animals’ recovery, the position of each electrode in the arrays was identified based on computer tomography, and then co-registered to a template T1-weighted anatomical magnetic resonance image (MRI) (http://brainatlas.brain.riken.jp/marmoset/; Hikishima et al., 2011) (monkey Fr) or pre-acquired MRI (monkeys Go and Kr) using MRIcron software (http://www.mricro.com; Rorden et al., 2007). In all monkeys, the electrode-array covered the frontal, parietal, occipital, and temporal cortices, including the primary auditory area (Fig. 1F–I).

**Figure 1.**
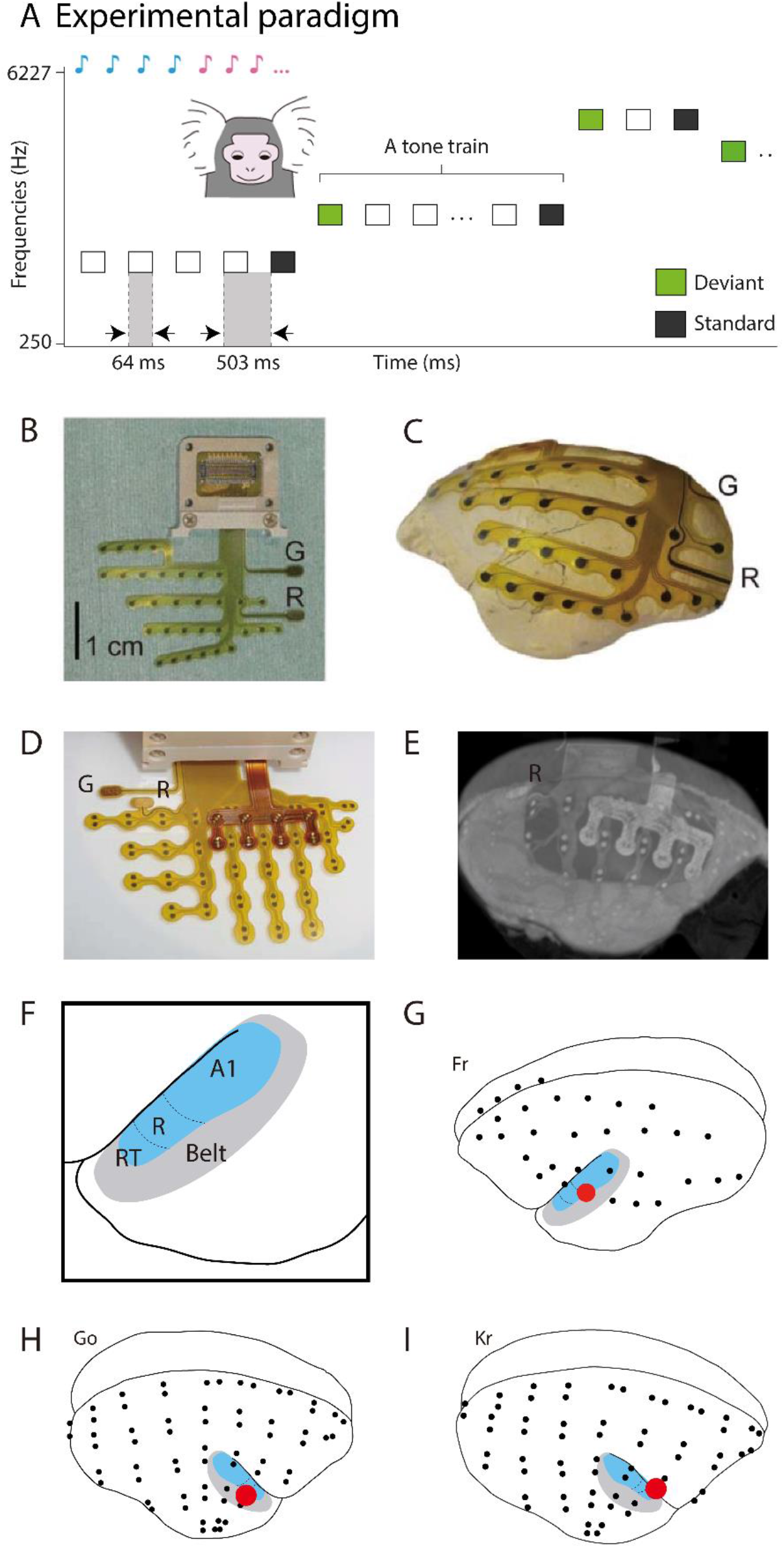
Experimental design and ECoG electrode arrays. **(A)** Using a roving oddball paradigm, 20 different single-tones were presented in the trains of 3, 5, or 11 identical stimuli. Any two subsequent trains consisted of different tones. This way, while the adjacent standard (depicted in black) and deviant (depicted in green) tones deviated in the frequency due to the transition between the trains, the two expectancy conditions were physically matched, as the first and the last tones of the same train were treated as deviant and standard tones in the analysis of the adjacent stimuli pairs. **(B)** The complete 32 electrode array and connector viewed from the front. G = ground electrode, R = reference electrode. **(C)** A fitting example of the 32 electrode array on a model brain. **(D)** The complete 64 electrode array and connector viewed from the front. **(E)** The CT image of the implanted 62 electrode array registered to the MRI of monkey Kr (two electrodes were cut during implantation). **(F)** An enlarged view of the temporal area of the marmoset, showing the core (A1, R, and RT) and belt areas. The borders between each auditory area are estimated by overlaying the Common Marmoset Brain Atlas (http://brainatlas.brain.riken.jp/marmoset/modules/xoonips/listitem.php?index_id=66) on the standard brain. A1, primary auditory cortex; R, area R (rostral auditory cortex); RT, area RT (rostrotemporal auditory cortex). **(G)** Locations of the 32 electrodes in the monkey Fr. The red circle indicates the electrode used for the spectral decomposition and MMA analyses. **(H)** Locations of the 64 electrodes in the monkey Go. The red circle indicates the electrode used for the spectral decomposition and MMA analyses. **(I)** Locations of the 62 electrodes in the monkey Kr. The red circle indicates the electrode used for the spectral decomposition and MMA analyses.

### Stimuli and task

We adopted a roving oddball paradigm (Cowan et al., 1993; Haenschel et al., 2005; Garrido et al., 2008). The trains of 3, 5, or 11 repetitive single-tones of 20 different frequencies (250–6727 Hz with intervals of 1/4 octave) were pseudo-randomly presented. Tones were identical within each tone-train but differed between tone-trains (see Fig. 1A). Because tone-trains followed on from one another continuously, the first tone of a train was considered to be an unexpected deviant tone, because it was of a different frequency from that of the preceding train. The final tone was considered to be an expected standard tone because it was preceded by several repetitions of this same tone. To avoid analytical artifacts stemming from differences in the number of standard and deviant stimuli, we considered only the last tone of a train as standard. There were 240 changes from standard to deviant tones in a single recording session. Pure sinusoidal tones lasted 64 ms (7 ms rise/fall), and stimulus onset asynchrony was 503 ms. Stimulus presentation was controlled by MATLAB (MathWorks Inc., Natick, MA, USA) using the Psychophysics Toolbox extensions (Brainard, 1997; Kleiner et al., 2007; Pelli, 1997). Tones were presented through two audio speakers (Fostex, Japan) with an average intensity of 60 dB SPL around the animal’s ear.

### ECoG recording and preprocessing

ECoG recordings were taken in the passive listening condition while monkeys were awake. In each recording session, the monkey Fr was held in a drawstring pouch, which was stabilized in a dark room, and the monkeys Go and Kr sat on a primate chair in a dimly lit room. The length of a single session was about 15 min: the first 3 min of data were used for another auditory paradigm consisting of standard tones (data are not shown in this paper) and the remaining 12 min of data were used for the roving oddball sequences. For monkey Fr, data from 3 sessions were used for analysis, which resulted in 720 (=240 × 3) standard and deviant presentations. For monkeys Go and Kr, data from 6 sessions were used for analysis, which resulted in 1440 (=240 × 6) standard and deviant presentations.

ECoG signals were recorded using a multi-channel data acquisition system (Cerebus, Blackrock Microsystems, Salt Lake City, UT, USA) with a band-pass filter of 0.3–500 Hz and then digitized at 1 kHz. In the signal preprocessing, those signals were re-referenced using an average reference montage, and high-pass filtered above 0.5 Hz, using a 6^th^ order Butterworth filter. We segmented datasets from −903 to 400 ms relative to the onset of the unexpected tone, so that each segment would include a pair of a deviant and a standard immediately preceding the deviant, as well as a baseline of 400 ms preceding the standard tone (see Fig. 1A). The segments were then further divided into standard epochs and deviant epochs (−400 ms to 400 ms with 0 ms indicating the onset of tone). While we intended to carry out the main analyses on shorter epochs in the present study (−100 ms to 350 ms, see below), we initially created wider epochs that were used for different analyses not reported here. For the spectral decoupling, ECoG segments of the longest 11-tone trains were created (−200 ms to 5500 ms with 0 ms indicating the onset of the first tone of the sequence (deviant tone), see below). Parts of the dataset are shared in the public server Neurotycho.org (http://neurotycho.org/; Nagasaka et al., 2011).

### Functional localization of electrodes-of-interest: Event-related potentials and time-frequency analysis

ECoG electrode-of-interest was identified functionally by contrasting LFP between standard and deviant stimuli (0-350 ms), separately for each electrode (see Fig. 2A). For functional localization of MMN waveforms, and only for it, a low-pass filter of 40 Hz was applied, using a 6^th^ order Butterworth filter. ECoG recordings were re-referenced with respect to the common average reference across all electrodes. Data were then epoched around the onset of tones (−100 ms to 350 ms), and baseline correction was applied by subtracting the mean of the 100 ms period before the stimulus onset. MMN was assessed by comparing the standard ERP and deviant ERP, and one electrode with the largest MMN amplitude was selected for each monkey for further analyses. In all three monkeys, the identified electrode-of-interest was located in the auditory cortex (see Fig. 1G–I). In addition, given that MMN is associated with high-gamma response (Dürschmid et al., 2016; Edwards et al., 2005; Eliades et al., 2014; Haenschel et al., 2000; MacLean et al., 2014; Marshall et al., 1996), we inspected whether the electrodes with peak MMN amplitude would also show the highest gamma power difference between standard and deviant tones. We plotted time-frequency charts between standard and deviant stimuli (0-350 ms), separately for each electrode. Epochs were first bandpass filtered in multiple successive 10 Hz wide frequency bins (from 1 to 250 Hz) using a zero phase shift noncausal finite impulse filter with 0.5 Hz roll-off. Next, for each bandpass filtered signal, we computed the power using standard Hilbert transform. Finally, the resulting time-frequency charts were z-score transformed from −100 to 0 ms. As expected, the largest gamma effect was indeed observed in the electrodes with the largest MMN amplitude (see Fig. 2A).

**Figure 2.**
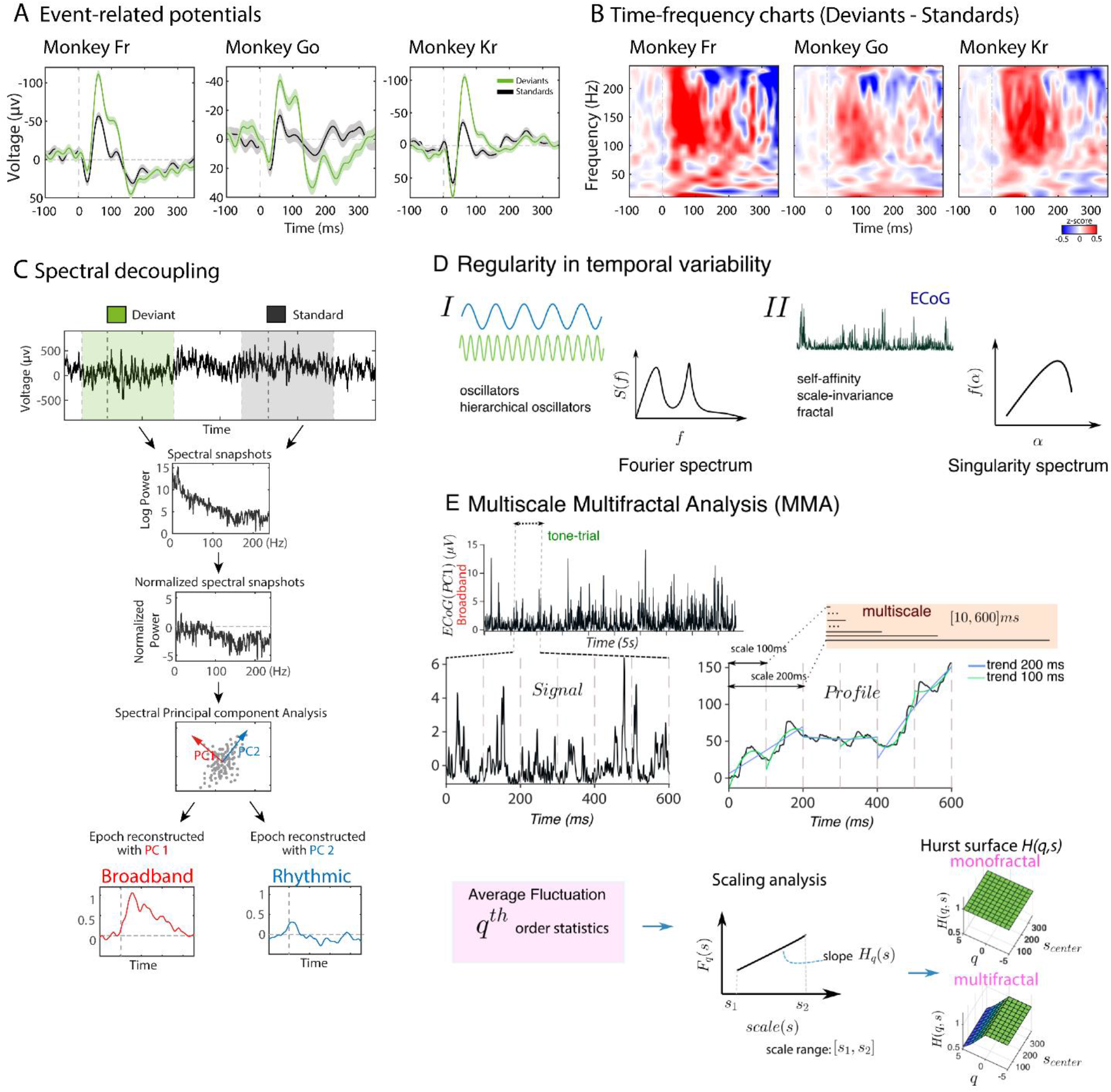
ECoG data analyses. **(A)** Time courses of ERP waveforms of the standard (black) and deviant (green) stimuli conditions. 0 ms time point indicates the onset of a given tone. Error shades represent the standard error of the mean (SEM), calculated across all trials at each time point. Each subplot represents a different monkey. **(B)** Time-frequency charts showing a spectral power response to the auditory stimuli, expressed as the difference between the standard and the deviant tones. The 0 ms time point indicates the onset of tones. **(C)** Spectral decoupling. Temporally adjacent raw LFP segments of the standard tone (i.e. the last stimulus of the previous train) and the deviant tone (i.e. the first stimulus of the subsequent train) were extracted for the spectral PCA. First, the Fast Fourier transform was used to calculate log power (1-250 Hz) of the raw LFP signal, which was afterwards normalized across all trials within a given expectancy condition. Normalized spectral snapshots were input into spectral PCA, which separated broadband and rhythmic components. Principal spectral components were reconstructed back to the time-series for the subsequent contrast between the expected and unexpected stimuli conditions. **(D)** Two views of regularity in temporal variability. Some simple machines display mechanistic behavior which can be described by decomposing their output variables into multiple frequency-dependent oscillators or combinations of these (spectral analysis). By contrast, brains display complex behavior with output variables which appear erratic, intermittent, and nonstationary and are the result of an inextricable interdependence of processes at many temporal scales. In this case, a measure that is scale-free and a “summary” of the whole activity is more adequate to characterize the regularities of the signal. This can be thought of as obtaining the spectrum of the different fractal/scaling exponents/singularities hidden in the signal (multifractal analysis). **(E)** Overview of Multiscale Multifractal Analysis (MMA) (cf. *Results, Methods*).

### Decoupling the cortical spectrum to isolate Broadband and Rhythmic spectral components

To extract the course of broadband spectral activity, we carried out the spectral decoupling of raw LFP signal (Miller et al. 2009a, 2009b, 2017; Miller, 2019). As for the ERP analysis, ECoG potentials were re-referenced with respect to the common average reference across all electrodes. For the selected electrodes-of-interest (see above), discrete epochs of power spectral density (PSD) were calculated using the long epochs corresponding to the 11-tone sequences (−200 ms to 5500 ms around the first tone (deviant tone) of each sequence). For each trial (i.e. each 11-tone sequence), individual PSDs were normalized with an element-wise division by the average power at each frequency, and the obtained values were log-transformed. The eigenvectors (Principal Spectral Components or PSCs) from this decomposition revealed distinct components of cortical processing, with the first three PSCs explaining most variance: the broadband PSC, i.e. no predominant peak present in the PSD, the first rhythmic PSC (alpha rhythm, ~ 10 Hz), and the second rhythmic PSC (delta rhythm, ~2 Hz) (see Fig. 2C and 3A,H,O).

**Figure 3.**
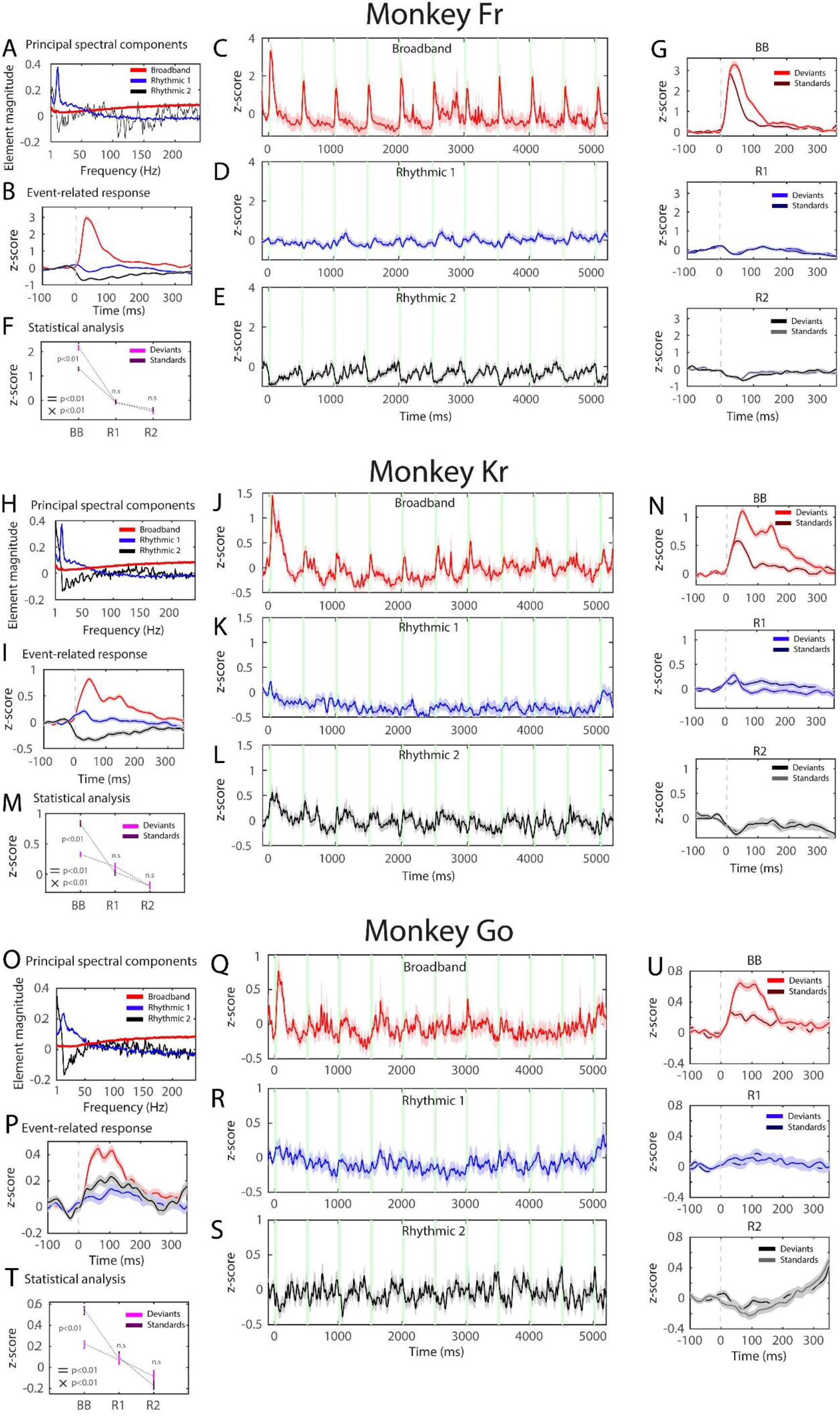
Event-related broadband response of stimulus expectancy. **(A,H,O)** Element magnitude of the major principal spectral components (PSCs) in the frequency domain (1-240 Hz). In this and other subplots, the Broadband PSC is depicted in red, the Rhythmic 1 PSC (alpha) in blue, and the Rhythmic 2 PSC (delta) in black. **(B,I,P)** A narrow window of back-reconstructed time series of the broadband and rhythmic PSCs, locked to the onset of tones (0 ms). Standard and deviant stimuli are averaged together. **(C-E,J-L,Q-S)** Back-reconstructed time series of the Broadband and Rhythmic PSCs along a sequence of 11 identical tones. 0 ms indicates the onset of the deviant tone. **(F,M,T)** ANOVA results of the stimulus expectancy (standard, deviant) and the spectral component (Broadband, Rhythmic 1, Rhythmic 2) contrast. Significant main effects were observed for the PSC (Fr: F(2,1438)=341.70, p<0.001, eta-squared = 0.322; Kr: F(2,2878)=113.00, p<0.001, eta-squared = 0.073; Go: F(2,2878)=78.60, p<0.001, eta-squared = 0.052) and the stimulus expectancy (Fr: F(1,719)=14.1, p<0.001, eta-squared = 0.01; Kr: F(1,1439)=23.60, p<0.001, eta-squared = 0.016; Go: F(1,1439)=4.81, p<0.029, eta-squared = 0.003) factors, and the interaction between the PSC and the stimulus expectancy (Fr: F(2,1438)=17.20, p<0.001, eta-squared = 0.02; Kr: F(2,2878)=20.80, p<0.001, eta-squared = 0.014; Go: F(2,2878)=15.49, p<0.001, eta-squared = 0.011). Error bars indicate the standard error of the mean (SEM). ‘=’ refers to the main effects, ‘x’ refers to the interaction. **(G,N,U)** Stimuli-locked waveforms show post-hoc comparisons between the standard and deviant stimuli in the broadband and rhythmic PSCs, which revealed larger amplitude for the deviant stimuli in the Broadband PSC contrast (Fr: t=6.96, p_Bc_<0.001; Kr: t=7.84, p_Bc_<0.001; Go: t=5.48, p_Bc_<0.001), but not in the Rhythmic 1 (Fr: t=0.378, p_Bc_=1.00; Kr: t=0.612, p_Bc_=1.00; Go: t=0.397, p_Bc_=0.99) nor the Rhythmic 2 (Fr: t=0.812, p_Bc_=1.00; Kr: t=−0.033, p_Bc_=1.00; Go: t=−1.567, p_Bc_=1.00) PSC contrasts. **Note:** For visualization purposes, 11-tone time series were smoothed with an 80-ms Gaussian envelope (SD 80 ms) and single-tone time series were smoothed with a 20-ms Gaussian envelope (SD 20 ms). However, all statistical analyses were carried out on non-smoothed data.

In order to identify components of stimulus-related changes in the PSD (Fig. 3), an inner product matrix of the normalized PSDs previously performed on the 11-tone sequences was diagonalized with a singular value decomposition, and was then applied to the individual epochs of −100 to 350 ms length with 0 ms indicating the stimuli onset for both standard and deviant tones (Fig. 3B,G,I,N,P,U); and to the 11-tone sequences themselves (Fig. 3C–E,J–L,Q–S). Following Miller et al. (2009), the entire time series were *z*-scored per trial (to put in intuitive units, because this measure is approximately normally distributed), exponentiated, and then a value of 1 was subtracted (setting the mean closer to 0). Then, in order to make both conditions comparable by setting the baseline period to 0, we further performed a baseline correction on the pre-stimulus period by subtracting the mean value per trial between −100 to 0 ms. The first PSC allowed us to obtain the “broadband time course” that has been shown to reflect a power law in the cortical PSD (Miller et al., 2009a), and the second and third PSCs uncovered the “rhythmic time courses” with distinct frequency peaks.

### Cross-individual decoding

To assess the generalizability of our findings in the temporal domain, we carried out between-subjects cross-decoding, known to be a more conservative method of validation compared to within-subjects decoding (e.g. Gevins et al., 1998; Visell & Shao, 2016). A univariate temporal decoding model was applied on each PSC time-courses on the selected auditory cortex electrodes, aiming to decode the stimuli expectancy categories, i.e. standards vs deviants (Fig. 4). The ADAM-toolbox was used on the Broadband and Rhythmic PSC time-courses with epochs from −100 ms to 350 ms (Fahrenfort et al., 2018). Crucially, and for each individual neural component, we trained a linear discriminant (LDA) classifier in one monkey and tested it in a separate monkey for obtaining cross-individual decodability of stimuli expectancy category, i.e. standard vs deviant trials. Next, a backward decoding algorithm, using either stimulus expectancy category was applied, training the classifier on one monkey and testing on another monkey. A linear discriminant analysis (LDA) was used to discriminate between stimulus classes (e.g. deviant versus standard trials) after which classification accuracy was computed as the area under the curve (AUC), a measure derived from Signal Detection Theory. This allowed us to determine whether standard could be discriminated from deviant better than chance. Instead of relying on a hypothetical chance level AUC, we nonparametrically computed the distribution of actual chance level accuracies for every time-point by running 500 iterations, randomly permuting the class labels on every iteration. This in turn allowed us to obtain a p-value for every time sample. Each *p*-value was computed by dividing the number of times that the AUC value under random permutation exceeded the actually observed AUC value, and dividing this value by the total number of random permutations. These nonparametrically obtained *p*-values were corrected for multiple comparisons over time, using a false discovery rate (FDR) correction (*p*<0.05, *q*=.05) (Benjamini & Hochberg, 1995; Benjamini & Yekutieli, 2001), thus yielding multiple comparisons corrected time windows of above-chance accuracy. This procedure allowed us to assess whether the decodability of one brain with a classifier trained on another would succeed at a meaningful time window of interest, i.e. around 50-100 ms when MMN was observed (see Fig. 2A). Notably, given that decoding was carried only on one electrode that showed the strongest effect, spatial heterogeneity between monkeys was removed entirely, which effectively left only one dimension, i.e. time, in which animals could display differences.

**Figure 4.**
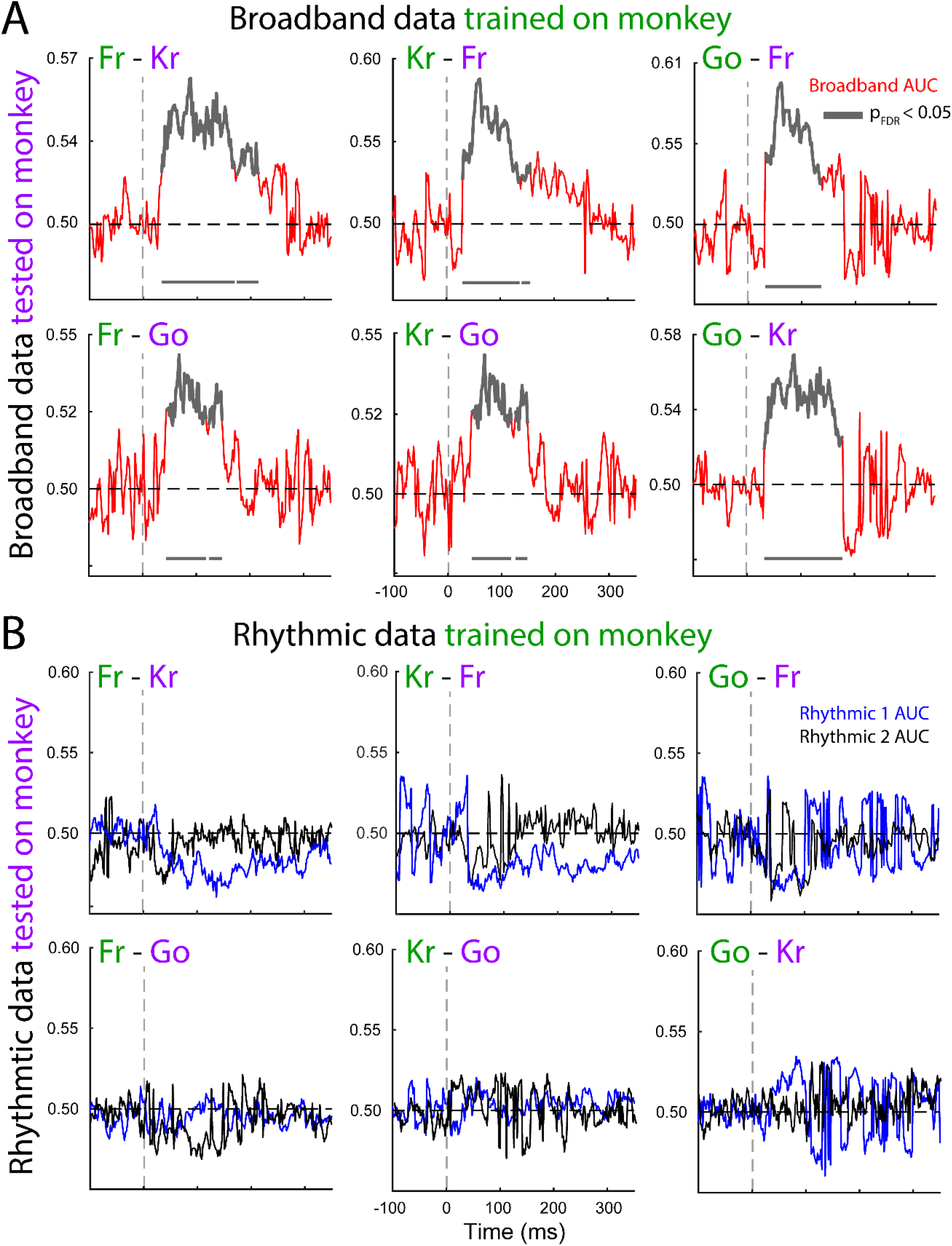
Decoding of stimulus expectancy across monkeys with Broadband and Rhythmic components. Classification of stimulus expectancy conditions (standard, deviant) was carried out in one of the monkeys (plotted in green). Afterward, the LDA classifier was tested on the other two monkeys (plotted in purple). Time points of FDR-corrected significant decoding (AUC) of stimulus categories above are depicted in grey (see Methods for details). **(A)** Decoding was successful in all six pairs using Broadband PSC. **(B)** Decoding did not exceed the chance level using the Rhythmic 1 and Rhythmic 2 PSCs.

### Dynamical characterization of the scaling behavior

To characterize the scaling properties of the neural activities of all PSCs of all monkeys we first quantified the relationship between log(power) and log(frequency) of all individual trials (trains of 11-tones of standards and deviants combined).

#### Continuous power spectral densities

The power spectral density (band: 1–100 Hz) of each trial (baseline removed) for the PSCs studied was computed by applying the modified Welch periodogram method as implemented in Matlab’s *pwelch()* function. We used 50% overlapping Hann windows of 1.024 s, 250 ms were removed from the beginning/end of each trial to avoid edge artifacts. Subsequently, we combined all individual trials for each PSC after removing their 200 ms baseline – the resulting series had a length of 1327920 (Monkey Fr) and 2655840 samples (Monkeys Kr and Go).

The functional repertoire of neural networks hinges on the multiscale temporal organization of their coordinated activity. We aimed at obtaining a “summary” metric of the macroscopic fluctuations of neural activity that could tease apart the dynamical behavior of the extracted ECoG components. The slope of the double logarithmic relationship between the frequency *f* and the power spectrum (*S(f)*) (Fig. 5A) – the spectral exponent (β) – can be conceived as such a measure. However, even though for, e.g., a linear composition of stationary oscillatory signals the power spectrum accurately represents the signal, for macroscopic neural activity which is inherently nonstationary and nonlinear (Paluš, 1996), spectral analysis can be ambiguous and may yield identical results to dynamically distinct processes (Stanley et al., 1999). A scaling analysis allows quantifying regularities in data with a wide-band spectrum – without being tied to the rigidity of an oscillatory/”clock-like” code – by using the concept of self-affinity (loosely self-similarity) (see also Fig. 2D). Self-affine signals are characterized by having a statistically similar structure at different levels of magnification, i.e. a level of persistence or “memory”. Persistence manifests in a signal as “random noise” superposed upon a background with many cycles. These cycles are, however, not periodic, i.e., they cannot be extrapolated with time (Mandelbrot, 1983). The degree of persistence is quantified by the scaling exponent, higher persistence signifies that the signal is less irregular at *all scales* than the ordinary Brownian motion (uncorrelated noise) (Mandelbrot, 1983). Thus, lower values of the exponent are synonymous with higher irregularity and unpredictability. Physiological signals often display different degrees of persistence depending on the range of timescales considered. In the PSC data, this was apparent from the spectrum analysis which revealed changing points (crossovers) in the decay of power with frequency, i.e., the decay was only piecewise linear, suggesting the PSC’s spectral densities were not purely self-affine. Furthermore, neural signals can be more heterogeneous. The degree of persistence may differ with the *moment* of the time series analyzed (Kantelhardt, 2011), i.e. while traditional scaling analysis quantifies the scaling relationship in a time series’ second-order statistics ((mono)fractality), a generalized analysis looks for scaling relationships in *q*th-order statistics, where *q* can take any arbitrary value. Quantifying the persistence of multiple interwoven fractal subsets (scalings) within the signal is commonly referred to as multifractal analysis. A signal is multifractal if it requires different scaling exponents, dependent on the order of statistics studied, for its description; conversely, the signal is monofractal if its scaling behavior is equal for all the moments analyzed (Fig. 2E-bottom).

**Figure 5.**
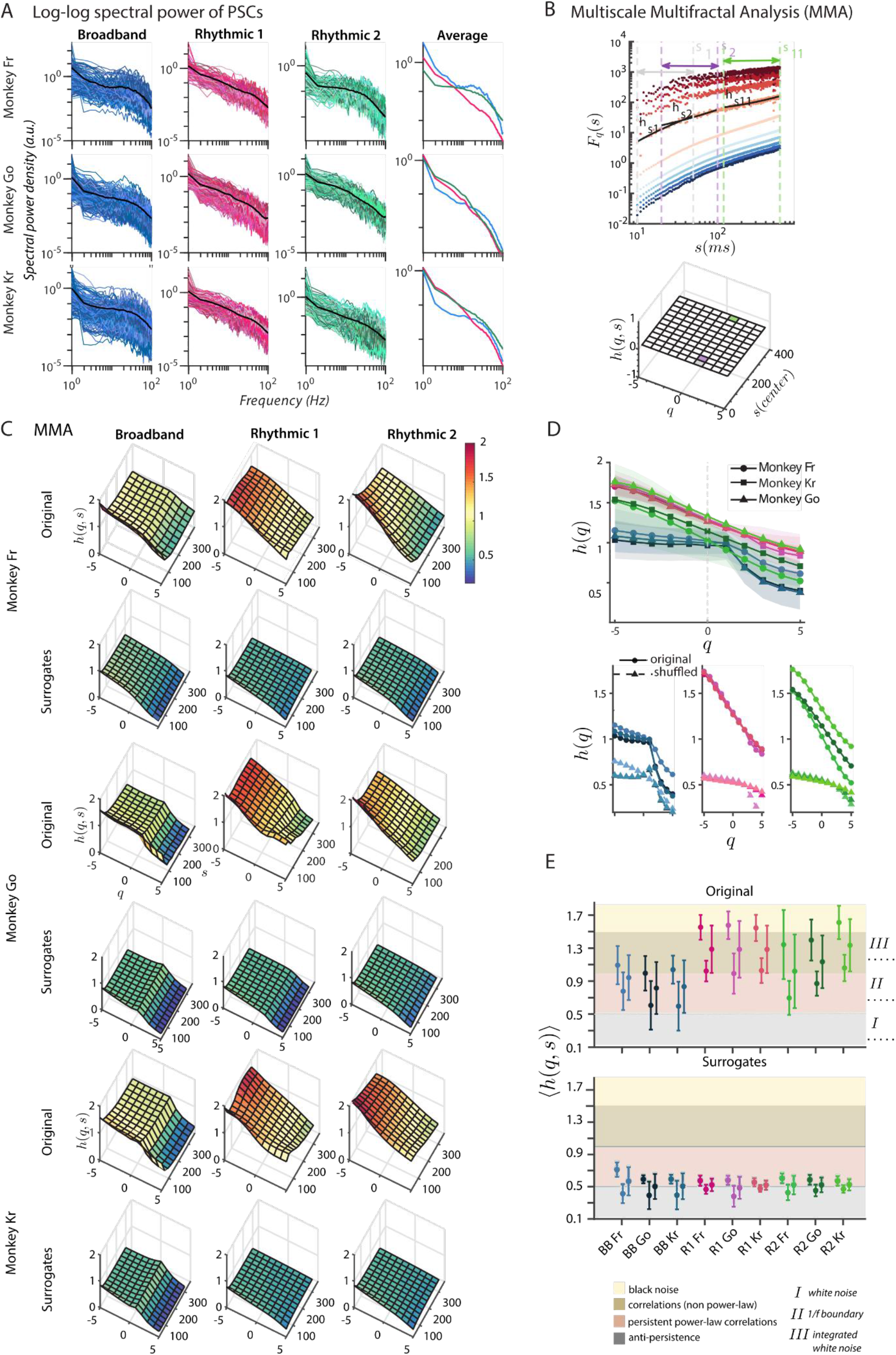
Multiscale multifractal analysis of the Broadband and Rhythmic dynamics. **(A)** Double logarithmic plots of the power spectral densities of the Broadband (blue), Rhythmic 1 (pink) and Rhythmic 2 (green) components during all trials (trains of standard and deviant tones) reveal a piecewise approximately linear decay of power with frequency. The average scaling (fractal) properties of the power spectral densities (last column and black lines in other columns) are distinct across frequencies, spectral components, and marmosets. Notably, while spectral power densities are plotted in the current figure, weights of spectral PCA across frequencies are plotted in Fig. 3A,H, O. **(B)** Multiscale Multifractal Analysis (MMA) method. *Top.* Log-log plots of the fluctuation functions *F*_*q*_(*s*) for each *q* ∈ [−5, 5], color-coded from dark blue (*q* = −5) to dark red (*q* = 5) and scale *s* (in ms) of the time series corresponding to the Broadband activity of monkey Go. The Hurst (scaling) exponent (*h*_1,2,…,11_) is obtained by determining the slope of a linear fit within a window lasting the period (*s*_1,2,…,11_) marked with vertical dashed lines. Three example scales are displayed: *s*_1_ ∈ [10, 50], and *s*_11_ ∈ [120, 600]. *Bottom.* Computed Hurst exponents *h*(*q*, *s*) are displayed in a (Hurst) surface plot grid. As an example, the cells corresponding to *h*_1,2,…,11_ (*q* = 2; *s* = 1, 2 or 11) are highlighted with their respective colors (light grey, lilac, green). **(C)** Hurst surfaces (*h*(*q, s*)) of the component activities (each column) and the < *h*(*q, s*) > of a distribution of 50 shuffled surrogates for each monkey. **(D)** Scaling properties averaged for all scales. The Hurst exponent *h* dependency on *q* is evident for all components, suggesting their multifractality. **(E)** Mean (+/-SD) of the Hurst surfaces (< *h*(*q, s*) >) suggests that the Broadband activity has an overall more irregular profile. Each group of 3 dots with error bars refers respectively to < *h*(*q, s*) > across all scales (*s*) for negative, positive, and all values of *q*. Individual results for the Broadband (BB), Rhythmic 1 (R1), and Rhythmic 2 (R2) PSCs are displayed in variations of blue, pink, and green colors, respectively. The bottom row shows the values obtained for the distribution of surrogates. The shaded colors denote the interpretation of *h(q,s) (relevant for panels C and D)* as the degree of persistence, the lower the value of *h*, the greater the irregularity or unpredictability in the signal.

#### Multiscale Multifractal Analysis

Since our goal was to study and contrast the multifractal properties of the different ECoG components and our preliminary spectral analysis revealed the location of crossovers (changes of scaling) varied across individuals and PSCs, a pre-definition of the timescales of interest was impossible. Thus, we used a method called Multiscale Multifractal Analysis (MMA) (Fig. 2E) (Gierałtowski et al., 2012), a data-driven scaling analysis robust to nonstationarity. MMA is an extension of the Detrended Fluctuation Analysis (DFA) (Peng et al., 1995), an established method to quantify the monofractal scaling behavior of nonstationary signals, robust to some extrinsic trends (Hu et al., 2001). DFA is essentially a modified root mean square (RMS) analysis of a random walk (Peng et al., 1995). Briefly, for a given time series *x*_*k*_ of length *N*, the profile *y(k)* is determined by integrating the time series, then *y(k)* is split into non-overlapping segments with length *s* which are detrended by subtracting the local least-squares line fit, *y*_*s*_(*k*). Since *N*/*s* is often not an integer, to avoid discarding data samples, a second splitting is performed starting from the end of the time series; a total of 2*N* segments are considered. The root-mean-square fluctuation of the integrated and detrended time series is given by:

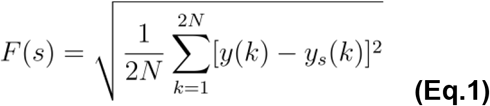

The Multifractal Detrended Fluctuation Analysis (MF-DFA) extension (Kantelhardt et al., 2002) permits to assess multifractality in the signals through generalizing the average fluctuation function *F(s)* of Eq. 1, in particular, the variance 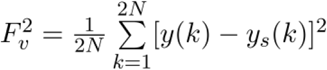:

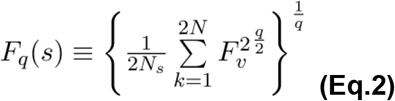

*Eq. 1* corresponds to the case *q* = *2* and assesses how the standard deviation changes with scale. This situation corresponds to the standard DFA, which is equivalent to the Hurst exponent (Hurst, 1951) for stationary random processes. By varying *q*, we can study shorter and longer fluctuations, i.e., how the *q-th* root of the *q-th* order variance changes with scale (*s*). Typically, *F*_*q*_(*s*) increases with *s* and displays the asymptotic behavior *F*(*s*) ~ *s*^*h(q)*^. The generalized Hurst exponent (*h*(*q*)) is estimated by extracting the slope of a linear least-square regression of *F*_*q*_(*s*) on a log-log plot for a given set of *s* values. The MMA algorithm allows scanning for several scale intervals yielding a quasi-continuous characterization of the scaling behavior (*h*(*q*)), which may vary among scales: the result is a scaling exponent depending on both *q* and *s*—*h*(*q, s*). It can be visualized in a grid – the *Hurst surface* – each grid cell corresponding to a value of *q* and a given interval of scales *s* (Fig. 2E, Fig. 5B).

We applied MMA to the PSCs of the 3 marmosets within a range of *q* ∈ [−5, 5], the negative values of *q* assess short fluctuations while the positive ones look at large fluctuations. The scale interval spanned 10-600 ms (~1.67-100 Hz), the former limit was chosen to avoid arithmetic underflow (Gierałtowski et al., 2012), and the latter to exclude scales beyond the length of a tone trial. We computed MMA along 12 scale intervals. The first scale integrated the scales *s*_1_ ∈ [10, 50] and for *s*_2,3,…,12_ this window was progressively slid 10 ms and expanded (*s*_2_ ∈ [20, 100], *s*_3_ ∈ [30, 150] and so forth). This permitted a nearly continuous coverage of the whole spectrum, allowing to identify crossover areas. For the detrending, we used a polynomial of order 2. The values of *h*(*q, s*) were interpreted in the following way (Gierałtowski et al., 2012): if *h*(*q, s*) = 0.5 the signal is constituted by uncorrelated randomness (white noise), *h*(*q, s*) ∈]0.5, 1] indicates persistent long-range correlations, if *h*(*q, s*) ∈ [0, 0.5[ the signal has anti-correlations, *h*(*q, s*) = 1 reveals 1/*f* (pink) noise and *h*(*q, s*) = 1.5 indicates Brownian motion (integrated white noise) and, finally, *h*(*q, s*) > 1.5 indicates black noise. Within the regime of *h*(*q, s*) ∈]0.5, 1], there is a straightforward correspondence between *h* and the spectral exponent *β* of the original (non-integrated) signal obtained from the slope of the power spectral density (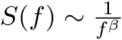, where *f* is the frequency); according to the Wiener-Khintchine theorem: *β* = 2*h* − 1. A full description of MMA is available at (Gierałtowski et al., 2012) and we used the original code available at Physionet (https://physionet.org/physiotools/mma/; Goldberger et al., 2000).

#### Multifractal and singularity spectrum

After obtaining *h(q,s),* we also computed the corresponding singularity spectrum. This representation yields a quantification of the different scaling relationships present (quantified by *h* above) paired with their respective proportion (density or dimension) in the signal. The scaling relationships are designated singularity strengths (*α*, the Lipschitz-Hölder exponent) and their densities *f*(*α*) (Halsey et al., 1986). Besides this generic goal, the motivation was to obtain indices that could summarize the differences in the multifractal properties of the neural components observed in the *Hurst surfaces* (elaborated below). This can be interpreted as an alternative way of thinking of spectrum to the classical Fourier spectral analysis (Fig. 2D).

One can connect the exponents obtained with MMA with the mass exponent *τ*(*q*), from the box-counting formalism in fractal geometry. This formalism allows studying the dimension of a given set *S* by relating, in a geometric support (such as a plane), a given distance *s* to the mass/population of points from the set. Considering that *S* has *N* points and *N*_*i*_ is the number of points in each box *i,* we can estimate the mass by the probabilities 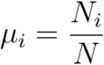. The *τ*(*q*) exponent is obtained through the relationship between the weighted number of boxes, *N(q,s),* and the moments of the probabilities, *μ*_*i*_ (Feder, 1988):

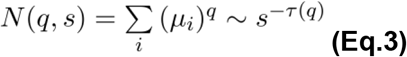

The exponent *τ*(*q*) varies with moment order (*q*) and characterizes how *μ*_*i*_ scales with *s,* it is often designated as the multifractal spectrum. This exponent relates to the *h(q)* exponents obtained with *MMA* by (Kantelhardt et al., 2002):

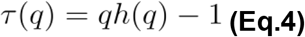

When *q* = 0, *τ*(0) = *D*_0_, which is referred to as the fractal dimension of the set. If there is a linear dependency of *τ*(*q*) with *q* then, a set is monofractal, otherwise, it is multifractal (Kantelhardt, 2011). It follows that *f*(*α*) is derived from *τ*(*q*) via a Legendre transformation (Halsey et al., 1986):

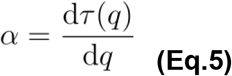

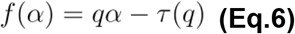

There are known caveats in the computation of *f*(*α*) with this method, namely its dependence on the differentiability of *τ*(*q*), but because of its compactness and consequent ease in extracting indices of multifractal properties, we opted to use it in consonance with MMA. For this reason, we recomputed the MMA values using still *q* ∈ [−5, 5] but with a step of 0.1 instead of 1, in order to obtain a smoother dependence of *τ*(*q*) on *q*. We verified that the functions *τ*(*q*) were reasonably smooth (Fig. 6B) so that *τ*(*q*) is probably everywhere differentiable.

**Figure 6.**
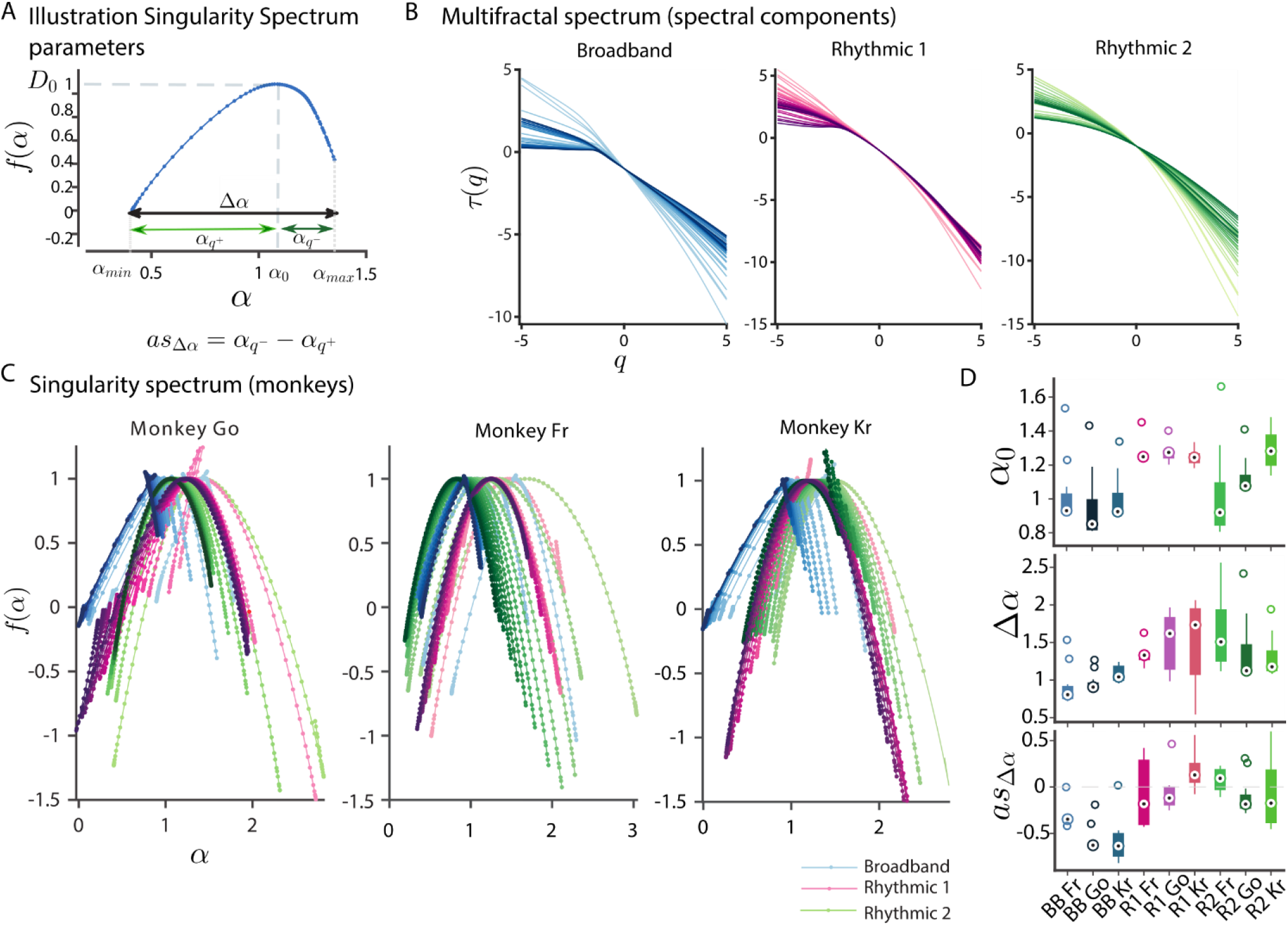
Contrast of the singularity spectra-based parameters of the Broadband and Rhythmic neural activity. **(A)** Illustration of the singularity spectrum (*f*(*α*)) and its parameters. *f*(*α*) reveals how densely the singularities (i.e., scaling exponents *α*) are distributed in a signal. The parabolic vertex shows the central tendency (*α*_0_, which corresponds to when *f*(*α*) = *D*_0_, the fractal dimension); the width Δ*α*, the degree of multifractality; the *α*_*q*_+ and the *α*_*q*_− represent the widths of the left and right tail, which correspond to values for *q* > 0 and *q* < 0. **(B)** Multifractal spectrum of the neural components; each graph shows a line for each scale range and monkey studied, the colors represent the different components (Broadband, Rhythmic 1 and 2) and the gradient light–>dark the scale range (s_1_, s_2_,…etc). Note that for the three components of neural activity, *τ*(*q*) is an approximately smooth function of q that is not linear, which reveals that signals are not monofractal (self-affine). **(C)** Singularity spectrum of the three PSCs for the three marmosets, the lightness of the colors represents the results for different scales (*s*) (light → dark with increasing scales *s*_1,2,… 11_). **(D)** Central tendency of the multifractal spectrum (*α*_0_), degree of multifractality (Δ*α*) and asymmetry of the spectrum (*as*_Δ*α*_) for the three types of activity ((Broadband (BB): blue; Rhythmic 1 (R1): pink; Rhythmic 2 (R2): green)). Each monkey is displayed in a different shade of the colors.

We aimed at obtaining several indices (for graphical overview, see Fig. 6A). It is noteworthy that the singularity spectrum typically has a convex upward shape and its left-branch (right-branch) corresponds to the *α*(*q*) values where *q* > 0 (*q* < 0) (Theiler, 1990). First, we obtained the singularity at the maximum of the curve (*α*_0_). At this singularity strength, the dimension corresponds to the fractal dimension (*D*_0_) of the measure supported by the set *S*, which is the union of all fractal subsets (*S*_*α*_) characterized by *α* in the continuum of *q* values. Since the complete set *S* has a dimension equal to *D*_0_, the subsets have fractal dimensions *f*(*α*) < *D*_0_ (Feder, 1988). Intuitively, multifractal analysis probes simultaneously the temporal structure of the aggregate (the singularity strength *α*_0_) and much finer structure of the time series (associated with the other singularity strengths) (Kelty-Stephen et al, 2013). Signals can have the same central tendency (similar *α*_0_) and, yet, interweave distinct combinations of singularity strengths. Next, to compare the degree of multifractality, we evaluated the width of the spectrum (Δ*α*). The Δ*α* is defined as the difference between the maximum (*α*_*max*_) and minimum (*α*_*min*_) values of the Lipschitz-Hölder exponent:

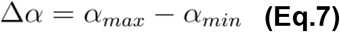

A narrow width implies the signal is monofractal (self-affine), the length of the width gives an indication of the deviation from monofractality, i.e., of the degree of multifractality (e.g., Ihlen, 2012). Remarkably, *f*(*α*) is not forcefully a symmetric function and can differ from the shape like the symbol characteristic of the most *trivial* multifractals, which are not strictly self-similar (scale-free), but have a multiplicative rescaling structure, i.e., a scale-dependent self-similarity (Riedi, 1999). Because of the above reason and the accentuated difference in the symmetry of *f*(*α*) between the broadband and rhythmic components, we quantified the following simple estimate of the degree of asymmetry:

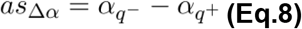

All parameters obtained from the singularity spectrum were computed for all marmosets and different timescales for each of the neural components and are presented individually or averaged across scales.

#### Surrogate data

We created 50 shuffled surrogates by randomly permuting in temporal order the samples of the original time series of each marmoset’s PSCs. If the shuffling procedure yields time series exhibiting simple random behavior (*h* = 0.5), one can conclude that the multifractality present is due to different long-range correlations of small and large fluctuations (Kantelhardt et al., 2002). On the contrary, if shuffling does not affect the values of *h*(*q, s*), the multifractality originates in a broad probability density function (PDF) of the values in the time series. If the multifractality originates both from correlations and broad PDF, the shuffling version will display weaker multifractality than the original one. All analyses were carried out in Matlab^®^ (v. 2018a, The MathWorks).

### Statistics

For ERP MMN (Fig. 2A) pairwise comparisons were used by comparing mean amplitude of pairs of adjacent standard (i.e. the last tone of the N train) and deviant (i.e. the first tone of the N+1 train) stimuli. Similarly, for the spectrally-decoupled time series (Fig. 3F,M,T), we performed separate repeated-measures ANOVA (RANOVA) of mean amplitude for each monkey between PSC (Broadband, Rhythmic 1, Rhythmic 2) and stimulus (standards, deviants), using Bonferroni correction for *post hoc* comparisons. Mean amplitude was always calculated over ±40 ms interval around the max peak detected after averaging all stimulus/component categories. In the case of the MMA (Fig. 5E), Hurst exponents were compared using RANOVA for each monkey between PSC (Broadband, Rhythmic 1 and Rhythmic 2) and *post hoc* comparisons were Bonferroni corrected. Statistical analyses were performed using open-source statistical software *jamovi* (Version 0.9; Jamovi project, 2019).

## Results

Using epidurally implanted electrodes, we recorded electrocorticograms (ECoGs) from three awake common marmosets, who passively listened to the stream of varying tones (see Fig. 1A). By contrasting neural responses to standard and deviant tones, which were physically matched across expectancy conditions, we first identified MMN deviance response from the auditory cortex (see Fig. 2A). Afterward, we decomposed the raw LFP signal into broadband and rhythmic spectral components (following Miller et al., 2009a, 2009b, 2017 and Miller, 2019; see Fig. 2C). Spectral decomposition allowed us to assess whether MMN is driven by the broadband rather than oscillatory components of the LFP signal. In the following, we report a single-trial analysis that was carried out separately for each monkey (referred to as Fr, Go, and Kr), using electrodes located in the auditory cortex (see Fig. 1G–I).

### Auditory ERP in the raw LFP signal

First, we confirmed that perturbation of auditory cortex with a deviant tone compared to a preceding standard tone increased the mean amplitude of auditory evoked potentials in the MMN time window (Fr: t(1,719) = −7.37, p < 0.001, Cohen’s d = 0.275; Go: t(1,1439) = −4.60, p < 0.001, Cohen’s d = 0.121; Kr: t(1,1439) = −9.27, p < 0.001, Cohen’s d = 0.244; see Fig. 2A). Latency of ERP peaks (58-66 ms) was consistent with the previous MMN studies of non-human primates (Javitt et al., 1992; Komatsu et al., 2015).

### Auditory evoked responses reconstructed with broadband and rhythmic components

Aiming to differentiate broadband component of LFP signal from rhythmic sources, we carried out spectral principal component analysis that decouples the power spectrum density (PSD) into components reflecting different underlying neural dynamics (see Fig. 2C and Methods). Using this technique, a broadband component can be identified by a uniform power increase, i.e. a component without clear peaks in the PSD, across a wide range of frequencies (see Fig. 3A,H,O, red lines). In addition to broadband spectral component, the technique also reveals a diverse set of narrow-band oscillatory components, revealed by peaks in the PSD (see Fig. 3A,H,O, blue and black lines).

This way, three major principal spectral components (PSCs), one representing a Broadband component and two representing Rhythmic components, were identified from the auditory LFP signal. PSCs were highly consistent across three monkeys (see Fig. 3A,H,O), matching tightly with the original depiction of spectral principal component analysis (PCA) (see Fig. 1A in Miller et al., 2009b). In order to assess which of these three major PSCs encode auditory prediction error response, components were back-projected to the time dimension.

We found that the Broadband PSC carried a characteristic auditory event-related broadband (ERBB) response, reminiscent of auditory ERP, compared to the relatively low amplitude of responses derived from the rhythmic PSCs with alpha (Rhythmic 1, ~9-11 Hz) and delta (Rhythmic 2, ~1-3 Hz) peaks. Arguably, the Rhythmic 1 component represents endogenous alpha oscillations generated in the auditory cortex (Billig et al., 2019; Haegens et al., 2005; Keitel and Gross, 2016), whereas the Rhythmic 2 component reflects temporal structure of exogenous stimulation, i.e. the stimulus-onset-asynchrony of 503 ms. The ERBB response was evident in the average of individual - standard and deviant - responses (see Fig. 3B, I, P) as well as along the whole sequence of 11 identical tones and a response was diminished in the tone sequences reconstructed from the Rhythmic components (see Fig. 3C–E,J–L,Q–S). Repeated measures ANOVA of mean amplitude between the PSC (Broadband, Rhythmic 1, Rhythmic 2) and the stimulus expectancy (standard, deviant) factors revealed the main effects for the PSC and the stimulus expectancy, and the interaction between the PSC and the stimulus expectancy factors (see Fig. 3F,M,T). Post-hoc comparisons showed that the ERBB response locked to the deviant tones had a larger amplitude compared to the ERBB response locked to the standard tones in the Broadband PSC contrast, but not in the Rhythmic 1 PSC nor the Rhythmic 2 PSC contrasts (see Fig. 3G,N,U). We thus conclude that MMN response recorded by the ECoG of the auditory cortex is driven by broadband rather than rhythmic components of the LFP signal.

### Cross-individual decoding of stimulus expectancy with broadband and rhythmic components

While the single-subject results of ERBB response were highly consistent across all three monkeys (see Fig. 3), we wanted to establish whether the broadband prediction-error response of an individual monkey can be extrapolated to other individuals of the same species. This would indicate that the prediction error information generated in the auditory cortex is implemented similarly across monkeys. We thus assessed the cross-individual generalizability of the ERBB response by decoding the stimuli expectancy using the Broadband and Rhythmic PSCs. Using all trials of a respective PSC of one monkey, we trained a linear discriminant (LDA) classifier to learn stimulus categories (standard vs. deviant) in the auditory cortex electrode (see Fig. 1G–I). Afterward, we decoded stimulus categories using the same PSC in a different monkey. Using Broadband PCS, we obtained significant decodability in all six pairs of comparisons, i.e. cross-individual decoding between 3 monkeys (see Fig. 4A). The time windows of significant decoding above chance level (50% AUC) were consistent with MMN and BRBB responses (see Fig. 2A and Fig. 3G,N,U). Contrary to this, no significant cross-individual decodability was observed using Rhythmic 1 and 2 PCSs (see Fig. 4B). These findings confirm the cross-individual generalizability of the broadband PSC encoding of stimulus expectancy.

### Multiscale multifractal analysis of broadband and rhythmic neural components

We hypothesized that during the evoked response, the multiscale temporal organization of the broadband differs from that of the rhythmic components. In particular, we sought to characterize and compare the scale-free temporal properties of the segregated neural components. These properties, both in terms of uniform scaling and intermittency (scale-dependent changes in scaling), are associated with perceptual and cognitive processing (Papo, 2014; He, 2014; Werner, 2010). We further hypothesized that the broadband component – the neural signal subserving oddball detection – has a more stochastic multiscale temporal organization which allows greater dynamical flexibility. The scale-free nature of the neuronal population firing rate, manifested in the broadband PSC (Miller, 2010; Manning et al., 2009), is usually estimated by determining the slope of the log-log function of PSD (power vs. frequency), also referred to as 1/*f* (fractal) scaling. However, often the PSD is not characterized by a single exponent and may show a scale-dependence (Miller, 2010; Chaudhuri et al, 2017) and/or different scaling depending on the statistical moment and hence exhibit multifractality (Nagy et al., 2017). Indeed, the single-trial auditory responses (standards and deviants) revealed a piecewise linear decay of power with frequency in each marmoset (Fig. 5A), suggesting that the dynamics of the underlying processes may have scale-free properties but also a heterogeneous scaling dependent on frequency (timescale). This is noticeable by the different slopes which characterize the 1/*f*-like PSD depending on the frequency range (Fig. 5A), precluding the fitting of a unique line to estimate the slope across the whole spectrum. Thus, to fully characterize the scale-free properties of the three components, we sought to test for the presence of scale-dependent multifractality in the series of increments of neural activity in the marmoset auditory cortex.

Multifractality requires the presence of different scaling exponents (*h*) of different moments of the fluctuations (*q*) over a wide range of timescales (*s*) (Kantelhardt et al., 2002). To quantify the multifractal behavior, we applied multiscale multifractal analysis (MMA; Gierałtowski et al., 2012) (see *Methods* and Figures 2E, 5B). In a nutshell, the raw fluctuations of the spectral components (Fig. 2C,E) were integrated (profile Fig. 2E) and split into windows of 10 to 600 ms (~ ISI Fig.1A), from which the quadratic trend (colored lines Fig.2E–-profile) was removed. The scaling exponent (*H*_*q*_*(s)*) characterizes the relationship between the average fluctuation of the integrated and detrended signal associated with a given *moment (q)* and the window of observation (scale–-*s*) (Figs. 2E, 5B). For *q* = *2*, standard deviation changes with scale are quantified, for *q* < 0, short-term variability and for *q* > 0, longer fluctuations (see Eq.2, *Methods*). *H*_*q*_*(s)* for a range of *q* and s values is represented by a surface, if the surface is ~flat the signal is monofractal, if *H*_*q*_*(s)* varies substantially with *q*, the signal is multifractal. Using MMA, we found that all the three PSCs show considerable variability in the values of the in their *h(q,s)* values *– Hurst surfaces*) (Fig. 5C). The Rhythmic components displayed similar surfaces, distinct from the nonlinear profile of *h* across *q* ∈ [−5, 5] of the Broadband PSC activity. These results were consistent across monkeys (Fig. 5C). The average tendency across scales revealed a nearly linear dependence of *h* with *q* for both Rhythmic components suggesting their underlying dynamics appear multifractal. Conversely, although the dynamics of the Broadband PSC are also multifractal (in the sense that its fractal properties depend on *q*), the profile is bimodal (different for small (*q* < 0) and large (*q* > 0) fluctuations (Fig. 5D)). We note that the conventional Hurst scaling analysis (*q* = 2 results) did not provide a clear distinction between the Broadband and Rhythmic 2 components. Furthermore, averaged surface values of *h* suggest the Broadband fluctuations can be quasi-stochastic (*h* ~ 0.5) or persistent without obeying strictly a power-law (*h* ~ 1.1), depending on whether large (*q* >0) or small fluctuations (*q* < 0) are considered (Fig. 5E). Conversely, Rhythmic 1 and Rhythmic 2 fluctuations ranged from being close to Brownian motion (integrated white noise, *h* ~ 1.5) to scale-free. There was a notable agreement on the values across monkeys (Fig. 5D,E). Thus, while all three PSC components showed scale-free properties, there were significant differences in the apparent stochasticity, expressed as *h(q)*, between the components (Go: RANOVA *F(2,20)*=103, *p*<0.001, eta-squared=0.339; Kr: RANOVA *F(2,20)*=134, *p*<0.001, eta-squared=0.404; Fr: RANOVA F(2,20)=40.2, *p*<0.001, eta-squared=0.228). For all three monkeys, the Broadband component exhibited lower *h(q)* values compared to the Rhythmic 1 (Go: *t*=−14.05, *p*<0.001; Kr: *t*=−13.39, *p*<0.001; Fr: *t*=−8.54, *p*<0.001) and Rhythmic 2 (Go: *t*=−9.54, *p*<0.001; Kr: *t*=−14.88, *p*<0.001; Fr: *t*=−6.64, *p*<0.001) components.

To determine whether the multifractality, depicted in the Hurst surfaces (Fig. 5C), is caused by the temporal correlations of the signal distribution, we created a distribution of shuffled surrogates, i.e. copies of the original data with identical mean, variance, and histogram distribution but no temporal structure. While the mean Hurst surfaces of the surrogates distribution showed for all monkeys a decrease in multifractality (*p*<0.001) (Fig. 5C), the averaged Hurst exponent values indicated that the neural dynamics approached randomness (*h = 0.5*) for all monkeys (Fig. 5E). Therefore, the multifractality is caused mostly by the temporal correlations but also by a fat-tailed probability distribution. We subsequently computed the multifractal spectrum, *f(α)*. Analogously to a Fourier analysis, i.e. the decomposition of a signal into a sum of components with fixed frequencies, *f(α)* can be understood as the decomposition of a signal into a set of exponents α (Mandelbrot, 2003) (Fig. 2D; Fig. 6A). Their relative presence in the signal is weighted by the *f(α)* function. The Broadband activity interweaved more densely sets of singularities that are less self-similar than those of the Rhythmic components and displayed a lower degree of multifractality and a more asymmetrical *f(α)* (Fig. 6C,D), suggesting its dynamics differs from simple multiplicative cascades. The shape of the multifractal spectra for the Broadband activity also displayed a right-truncation (Fig. 6C), which is expected due to the leveling of *h(q,s)* for *q* < 0 (Ilhen, 2012).

To sum up, MMA revealed that the generalized scale-dependent Hurst exponent *h(q,s)* and the derived *f(α)* curves of the dynamics of Broadband and Rhythmic components show multifractality as well as marked differences of this property. Importantly, the Broadband component more closely approached a stochastic asymmetrical multifractal distribution.

## Discussion

In the present study, we compared two alternative views of prediction error processing, namely whether LFP oscillatory vs. broadband components of neural activity encode deviant sensory stimuli. We first replicated previous research by showing that auditory MMN peaks in the auditory cortex. Afterward, we separated the LFP signal into Broadband and Rhythmic components and repeated MMN analyses separately for each component. While the main two Rhythmic components present in the data were not able to distinguish between the standard and deviant tones, the Broadband component indexed the stimuli difference in the auditory cortex, confirming our first hypothesis. The findings were highly consistent across all three marmosets, and the cross-individual decoding in the temporal domain successfully classified stimuli category (standard or deviant) when data were trained on one monkey and tested on a different one. While the decoding accuracy was modest, i.e. AUC values ranged between 0.52 and 0.60, cross-individual decoding is more conservative due to individual differences such as the curvature of cortical gyri. Importantly, significant decoding was observed only with the Broadband PSC, as the decoding was unsuccessful with the Rhythmic PSCs. Our findings suggest that auditory MMN response, a classical marker of prediction error, is primarily driven by the broadband component of LFP signal.

Contrary to the oscillatory narrow-band signals, such as alpha rhythm, broadband activity recorded with ECoG primarily reflects asynchronous neuronal firing (Buzsáki et al., 2012; Guyon et al., 2021; Henrie and Shapley, 2005; Hwang and Andersen, 2011; Li et al., 2019; Mukamel et al., 2005). Likewise, our earlier modeling of power-law changes in Broadband PCS suggests they reflect a change in mean population-averaged firing rate (Miller et al., 2009a; Miller, 2010). The association between broadband signals and neuronal spikes has been demonstrated both during passive tasks (Guyon et al., 2021; Manning et al., 2009) as well as in response to sensory stimulation (Henrie and Shapley, 2005; Hwang and Andersen, 2011; Li et al., 2019; Medvedev and Kanwal, 2004). While most of the evidence comes from visual neuroscience (e.g. Henrie and Shapley, 2005; Hwang and Andersen, 2011), a direct link between broadband activity and neuronal firing has been also demonstrated in auditory cortices (Medvedev and Kanwal, 2004; Mukamel et al., 2005), although late broadband high-frequency response in A1 may also reflect dendritic processes separable from action potentials (Leszczyński et al., 2020). Overall, given that broadband PSC primarily reflects the mean firing rate of neuronal populations, and that neuronal spiking correlates with the high-frequency LFP in the auditory cortex, including post-stimulus responses, our findings suggest that prediction error responses are encoded through more flexible, dynamical, unstable broadband codes than relatively more stable oscillatory codes in the post-stimulus time window.

While it has been argued that a stimulus-driven phase reset of slow frequencies in the range of delta and theta oscillations may underlie prediction error response (Arnal et al, 2015; Fuentemilla et al., 2008; Ko et al., 2012), we found that the Rhythmic 2 component with a distinctive delta peak and a considerable contribution from theta range activity (Fig. 3A,H,O) did not discriminate between standard and deviant tones. Likewise, the Rhythmic 1 component representing alpha range activity did not encode prediction error response, even though several previous studies linked MMN to alpha band power (Ko et al., 2012; MacLean et al., 2014). Yet, despite of these discrepancies, our findings do not imply that low-frequency counterparts of neural activity do not contribute to predictive coding: long-term dependencies are relevant in sensory prediction in the auditory cortex (Rubin et al., 2016), and low-frequency neural oscillations are instrumental for enhancing (Schroeder and Lakatos, 2009) and gating information in the auditory cortex (Lakatos et al., 2013). In particular, given that we did not separate evoked and induced neural activity, it is possible that low-frequency ongoing oscillations could constrain predictive error responses during the pre-stimulus period. As our analyses focused solely on the post-stimulus responses, follow-up studies should investigate the role of Broadband and Rhythmic PSCs in the evoked vs. induced components of neural signals at both pre-stimulus and post-stimulus time windows.

Confirming the second hypothesis, our results demonstrate that prediction error processing is subsumed by an asynchronous broadband activity with dynamical properties very distinct from that of the rhythmic components. Importantly, this difference is unveiled when a multiscale approach is used to characterize fluctuations with several degrees of resolution (multiple fractal hierarchies) and it is patent in the surfaces and multifractal spectrum; the difference is equivocal by simply observing the power spectral densities or doing a classical Hurst analysis. The broadband component is distinctive from the other components by its lower level of self-similarity and multifractality and also by its asymmetric multifractal spectrum. Importantly, all the broadband and rhythmic components displayed multifractal fluctuations. We interpret this finding as multifractality reflecting a generic feature of neuronal networks, with cognition operating concurrently with modulations of this property (Papo, 2014). Arguably, spike trains represent information with a multifractal temporal coding (Fetterhoff et al., 2015) and the integrated multifractal spectrum permits to infer the tuning curve of spiking activity in primates (Fayyaz et al, 2019). This could be a more effective dynamical fingerprint of how information is encoded in neuronal assemblies than the one provided by oscillatory rhythms. This hypothesis is bolstered by ideas that synchronization *per se* only arises in collective states where no new information can be created. In contrast, adaptive behavior emerges from more subtle forms of coordination, e.g. through the metastability or asynchronous coupling of spatiotemporal patterns of neural activity (Friston, 2000; Tognoli and Kelso, 2014). The multifractality present in the recordings reveals how the macroscopic neural dynamics is intermittent, its spectral density changes with time, which has been hypothesized as a facet of temporal metastability (Friston, 1997; Tognoli and Kelso, 2014); at the core of metastability is the broken symmetry of spatiotemporal patterns (Kelso, 1995) which was only present in the broadband activity. Thus, it is perhaps this facet of multifractality that is relevant. In fact, the more asymmetrical multifractal spectra of the broadband activity suggest this feature may be a proxy of a dynamical regime that allows the breakdown of symmetry, characteristic of systems that can perceptibly or meaningfully react to afferent inputs (Freeman and Vitiello, 2006).

Furthermore, the prediction error processing by neural assemblies in the auditory cortex is sustained by an irregular broadband component with certain dynamical behavior. Specifically, tone and after-tone processing is characterized by small fluctuations lying in a tight range of the non-ergodic dynamical regime (*h* just above 1) – which has been proposed as an explanation for the 1/*f* noise of cognitive processes (Grigolini et al., 2009) – and large fluctuations with apparent weak correlations. The latter finding aligns with sensory discrimination of visual stimuli by monkeys being more accurate if network activity in the inter-stimulus period is more desynchronized (Beaman et al., 2017). Moreover, it is substantiated by theoretical accounts that used simulated recurrent neural networks to show that correlated presynaptic input and weakly correlated network spiking activity can coexist (Renart et al., 2010) and that asynchronous irregular states are responsive to afferent stimuli while synchronized oscillatory states are not (Zerlaut and Destexhe, 2017). Arguably, the arrhythmic components of the signal, while often discounted as ‘noise’, enable neural dynamics to be ready to jump from one state to another quickly. In such a case, the background neural ‘noise’ can continuously reorganize itself by staying in an active predictive state that is constantly ready to change depending on the incoming stimuli and internal prediction error responses.

Our findings were enabled by a novel approach to quantify these complex dynamics of neural systems, the so-called brain’s “stochastic chaos” (Freeman et al., 2001). Future studies are anticipated to extend MMA to wider frequency ranges (>100 Hz), with a fine-grained resolution to arguably uncover the spike tuning underlying sensory-state discrimination (Fayyaz et al., 2019). The broadband prediction error response should be further studied using hierarchical auditory prediction paradigms that can discriminate sensory and top-down prediction error responses (Bekinschtein et al., 2009; Chennu et al., 2013, 2016). Developed in human studies, such paradigms have been recently successfully applied in the common marmosets (Chao et al., 2018). Furthermore, while the marmoset model of MMN is deemed successful and very stable, as indicated by cross-individual decoding, the current study should be replicated using LFP recordings in humans.

Importantly, our findings reveal a meso-scale neural dynamics of MMN and contribute to a unifying framework for the micro- to macro-level neural mechanisms of the prediction error response. While most of the auditory MMN studies are carried out at the macro-level using scalp EEG recordings or meso-level LFP, auditory prediction error responses have also been identified using single-neuron recordings (Nieto-Diego and Malmierca, 2016; Parras et al., 2017; Pérez-González et al., 2005; Solomon and Kohn, 2014; Ulanovsky et al., 2003, 2004). In particular, individual neurons located in the primary auditory cortex increase spiking rate following presentation of oddball stimuli, which has been observed in different mammal species, including cat (Ulanovsky et al., 2003, 2004), rat and mouse (Nieto-Diego and Malmierca, 2016; Parras et al., 2017). Similar responses have also been identified in sub-cortical neurons (Parras et al., 2017; Pérez-González et al., 2005). In particular, a subclass of neurons located in the dorsal and external cortices of the inferior colliculus of the rat respond selectively to novel auditory stimuli, while muting their response to repetitive stimuli (Pérez-González et al., 2005). A recent study of single-neuron activity recorded from different auditory centers in rats and mice suggests that prediction error response is organized hierarchically along the non-lemniscal auditory pathway comprising of inferior colliculi, medial geniculate bodies and the primary auditory cortex with sensitivity to the deviant tones increasing along the pathway (Parras et al., 2017). MMN-like deviance sensitivity of firing rate increases further in the non-primary regions of the auditory cortex (Nieto-Diego and Malmierca, 2016). How do such micro-level single-neuron responses relate to the MMN potentials recorded with ECoG and/or EEG? Are there different neuronal mechanisms at different levels of measurement, such as single neuron spiking rate vs. neuronal oscillations recorded using ECog/EEG?

Our study indicates that increased neuronal firing rate may underlie prediction error responses not only at the micro-level of single-neuron recordings, but also at the higher meso-level LFP measurements. In particular, we show that MMN prediction error response is driven by the Broadband component of the meso-level LFP signal. Given that the Broadband PSC reflects largely stochastic neuronal firing rate, as suggested by previous modelling studies (Miller et al., 2009a; Miller, 2010), our findings are consistent with the account that auditory prediction error response is indeed encoded at a single action potential level within neuronal populations, which generate broadband signal at the meso- and most likely macro-level electrophysiology. Broadband LFP activity provides indirect access to the total spiking output of neurons, as shown by a growing number of experiments and simulations (Crone et al., 2011; Freeman, 2004; Rash et al., 2008). Thus, the reported Broadband activity in this study provides a ‘proxy’ for investigating the neuronal mechanisms underlying auditory prediction error. As such, the mesoscopic information of the Broadband LFP component represents a crucial link between macroscopic-level EEG and the microscopic-level spiking activity of neural populations (Buzsaki et al., 2012).

How could our LFP-based broadband results be reconciled with abundant literature on frequency-specific MMN results, mostly derived from EEG experiments that do not find broadband MMN response across all frequencies? First, the low-frequency range of broadband effects can be obscured by coincident changes in specific rhythmic phenomena (Miller, 2010). Second, too often classical frequency bands are loosely equated to specific rhythms (Lopes da Silva, 2013) and the views of collective neural network activity as oscillations lend too much emphasis on “rhythmicity” (Cole and Voytek, 2017) when in reality, in those narrow-band analyses perhaps no characteristic frequency oscillation was present and/or may even be spurious and caused by filtering (de Cheveigné and Nelken, 2017). Third, EEG artifacts may decrease signal-to-noise ratio in certain segments of the broadband signal, in which case only relatively clean segments would survive as significant detectors of prediction error response, especially when the effect sizes are small. For instance, blink artifacts may distort neural signals in the delta and theta frequency range (Gasser et al., 1992), whereas muscular artifacts are likely to interfere with the beta and gamma range activity (van de Velde et al., 1998). Given that unexpected stimuli modulate blinking rate (Bonneh et al., 2015) and facial expression (Reisenzein & Studtmann, 2007), an extreme case of which is a startle reflex (Brown et al., 1991), different levels of ocular and muscular EEG artifacts would be expected in response to standard and deviant tones. Furthermore, MMN amplitude and latency are modulated by drowsiness (Nashida et al., 2000; Sallinen and Lyytinen, 1997; Winter et al., 1995). Given that spontaneous fluctuations of alertness level are likely to occur during passive “oddball” paradigms, episodes of increased drowsiness could interfere with neural processing in the theta and alpha frequency range (Noreika et al., 2020a, 2020b). Thus, depending on the experimental demands, the selection and training of participants, and the data preprocessing steps, certain segments of the broadband signal may be occluded by artifactual or irrelevant signals when contrasting standard and novel stimuli, yielding narrow-band deviance responses that in fact originate from scale-free broadband component of the neuronal signal. The speculative role of EEG artifacts in the preclusion of broadband response should be tested in the future using simultaneous EEG and LFP recordings.

To conclude, we show that in a well-studied paradigm of auditory oddballs, oscillations do not constitute a means to temporally constrain information processing of auditory prediction error. They are perhaps the tips of the iceberg, the latter being an arrhythmic broadband component with asymmetric multifractal stochastic properties at several timescales. Our paper establishes the relevance of the broadband activity to encode relatively low-level auditory patterns and provides a theoretical background and empirical tools to probe which predictive values lie under the “noisy” surface in other paradigms and sensory modalities. While the present study focused exclusively on the local prediction error response in the auditory cortex, follow-up studies should investigate whether broadband or oscillatory signals underlie the top-down fronto-temporal predictions of incoming stimuli (Chennu et al., 2016; Garrido et al., 2008; Wacongne et al., 2011).

## Acknowledgments

We thank Dr Daniel Bor, Dr Laura Imperatori, Dr Robin Ince and Viktor Sadílek for contributing to valuable discussion, and Prof. Karl J. Friston for helping to sharpen theoretical aspects of the study. This manuscript is dedicated to the memory of Prof. Walter J. Freeman (1927 - 2016) whose pioneering work on Neurodynamics has inspired countless meaningful insights during the execution of this project.

## Notes

http://neurotycho.org/

### Competing Interest Statement

The authors have declared no competing interest.

